# Narcolepsy and the Dissociation of REM Sleep and Cataplexy through Ambient Temperature Manipulation

**DOI:** 10.1101/2021.12.29.474449

**Authors:** Bianca Viberti, Lisa Branca, Simone Bellini, Claudio LA Bassetti, Antoine Adamantidis, Markus Schmidt

## Abstract

Narcolepsy is characterized by increased REM sleep propensity and cataplexy. Although narcolepsy is caused by the selective loss or dysfunction of hypocretin (Hcrt) neurons within the lateral hypothalamus (LH), mechanisms underlying REM sleep propensity and cataplexy remain to be elucidated. We have recently shown that wild type (WT) mice increase REM sleep expression when exposed to thermoneutral ambient temperature (Ta) warming during the light (inactive) phase. We hypothesized that the loss of Hcrt may lead to exaggerated responses with respect to increased REM sleep and cataplexy during Ta warming. To test this hypothesis, Hcrt-KO mice were implanted for chronic sleep recordings and housed in a temperature-controlled cabinet. Sleep-wake expression and both spontaneous cataplexy and food-elicited cataplexy were evaluated at constant Ta and during a Ta manipulation protocol. Here we show several unexpected findings. First, Hcrt-KO mice show opposite circadian patterns with respect to REM sleep responsiveness to thermoneutral Ta warming compared to WT mice. As previous demonstrated, WT mice increased REM sleep when Ta warming is presented during the inactive (light) phase, whereas Hcrt-KO showed a significant decrease in REM sleep expression. In contrast, Hcrt-KO mice increased REM sleep expression upon exposure to Ta warming when presented during the active (dark) phase, a circadian time when WT mice showed no significant changes in REM sleep as a function of Ta. Second, we found that REM sleep and cataplexy can be dissociated through Ta manipulation. Specifically, although Ta warming significantly increased REM sleep expression in Hcrt-KO mice during the active phase, cataplexy bout number and total cataplexy duration significantly decreased. In contrast, cataplexy expression was favoured during Ta cooling when REM sleep expression significantly decreased. Finally, video actigraphy and sleep-wake recordings in Hcrt-KO mice demonstrated that Ta manipulation did not significantly alter waking motor activity patterns or waking or NREM sleep durations. These data suggest that neural circuits gating REM sleep and cataplexy expression can be dissociated with Ta manipulation.

**Statement of Significance:** Cataplexy and the loss of muscle tone has historically been viewed as a component of REM sleep that inappropriately intrudes into wakefulness. In addition to fragmented sleep, the intrusion of REM-like events into wakefulness has led to the hypothesis that narcolepsy represents a dysregulation of boundary state control. We show that REM sleep and cataplexy can be dissociated during the dark (active) phase through Ta manipulation. Such dissociation may provide clues regarding the cause of boundary state instability in Hcrt-KO mice, as well as provide a new method to interrogate mechanisms of REM sleep and cataplexy.

## Introduction

Narcolepsy with cataplexy (narcolepsy type 1) is a chronic sleep disorder affecting approximately 1 in 2,000 individuals and is caused by the selective loss or dysfunction of hypocretin/orexin (Hcrt) neurons in the lateral hypothalamus (LH)^1–4^. Narcolepsy is characterized by excessive daytime sleepiness (EDS), hypnopompic and hypnagogic hallucinations, sleep paralysis, sleep-wake instability and cataplexy. Cataplexy is the sudden loss of skeletal muscle tone triggered by generally strong and positive emotions during a state of preserved consciousness. Both cataplexy and rapid eye movement (REM) sleep share several common features such as a generalized skeletal muscle atonia, but it is still under debate whether cataplexy should be considered as an intrusion of REM sleep elements into wakefulness or a distinct state of brain activity^5–7^.

An intriguing aspect of narcolepsy is that several conditions are known to dissociate REM sleep and cataplexy. This dissociation is most evident as a circadian effect where REM sleep expression is highest during the circadian inactive (light) phase in mice at a time when cataplexy is rarely observed, whereas cataplexy is generally observed during the circadian active (dark) phase when REM sleep is reduced. Additionally, selective REM sleep deprivation over two consecutive days in humans with narcolepsy Type 1 leads to an increase in REM sleep pressure but without an increase in cataplexy^8^. Moreover, some medications such as selective D2/D3 antagonists have been shown to decrease cataplexy without affecting REM sleep expression^9^. These and other data, including the finding of differing EEG patterns between REM sleep and narcolepsy^6^, suggest that neural mechanisms triggering these two events may differ. However, the neural mechanisms underlying the ability to dissociate REM sleep and cataplexy remain unknown.

Another defining feature of narcolepsy is an increased REM sleep propensity^3,4^. For instance, patients with narcolepsy often show direct transitions from wakefulness into REM sleep known as sleep onset REM periods (SOREMs). Indeed, the multiple sleep latency test (MSLT) is utilized as a diagnostic clinical tool where patients with narcolepsy typically show two or more naps with SOREMs or a markedly reduced REM onset latency of 15 minutes or less in the nocturnal polysomnogram. A similar increase in REM sleep propensity has been demonstrated in mice lacking Hcrt neurons or receptors, characterized by increased REM sleep expression during their active wake phase^1,10,11^.

Although the Hcrt system may suppress REM sleep during waking^12,13^, the cause of both increased REM sleep propensity and cataplexy in narcolepsy has remained unclear. Recent data suggest that an overactive melanin-concentrating hormone (MCH) system may play such a candidate role. Indeed, using genetically modified hypocretin knockout (Hcrt-KO) mice to specifically target MCH neurons, recent work shows that chemogenetic activation of the MCH system increases both REM sleep and cataplexy, whereas MCH antagonism reverses this effect^14^. However, another study using calcium imaging found contradictory results, showing that MCH neurons are indeed active in REM sleep but silent during cataplexy^15^. Moreover, MCH-Hcrt double ablated mice show a marked increase in cataplexy and decrease in REM sleep compared to selective Hcrt ablated animals^16^. These latter data would suggest that although the MCH system may play a role in REM sleep expression, its inactivity would favor cataplexy.

Hcrt and MCH neurons are anatomically intermingled within the lateral hypothalamus and exhibit wide-spread projections to similar brain areas^17,18^. However, Hcrt and MCH neurons display reciprocal firing patterns^19^. For example, MCH neurons are active during REM sleep and their optogenetic stimulation enhances REM sleep transitions and prolongs REM sleep bout durations^20,21^. Hcrt neurons, in contrast, are predominantly active in wake and facilitate sleep to wake transitions^22–24^.

We hypothesized that the loss of Hcrt function in narcolepsy may lead to a disinhibition of the MCH system, potentially driving increased REM sleep propensity or cataplexy. Our prior work demonstrates a key role for the MCH system in driving REM sleep expression during thermoneutral ambient temperature (Ta) warming^25^. We used this thermoneutral Ta manipulation protocol to determine whether Hcrt-KO mice show exaggerated responses in REM sleep propensity and cataplexy expression as a function of Ta. We found that wild type (WT) and Hcrt-KO mice show opposite circadian responsiveness patterns with respect to REM sleep expression in response to Ta warming. Moreover, we demonstrate that although thermoneutral Ta warming increased REM sleep expression in Hcrt-KO mice during the active (dark) circadian phase, these mice markedly and significantly decreased cataplexy expression independent of waking activity during Ta warming. These data demonstrate a previously unknown ability of Ta manipulation to dissociate REM sleep and cataplexy in narcolepsy.

## Materials and Methods

### Mice

Two genetically-modified mouse lines were used for these experiments, including Hypocretin-knockout mice (Hcrt-KO) and MCH receptor-1-knockout (MCHR1-KO) mice. Homozygous Hcrt-KO (n=11), MCHR1-KO (n=11), and C57BL/6 WT (n=22) male mice were used for these experiments using methodology as previously described^25^. The genotyping for all mice was verified using PCR from ear clip biopsies. Mice were aged between 8-20 weeks.

Experiments were performed at the Zentrum fur Experimentelle Neurologie (ZEN) at the University Hospital, Inselspital, in Bern, Switzerland. All the experiments were carried out in accordance with the guidelines described in the National Institutes of Health Guide for the Care and Use of Laboratory Animals and the Bern Kanton Animal Care Committee.

### Surgical procedure for electrophysiological recordings

Animals were anesthetized with isoflurane (2% in O2) and injected with Metacam (0.1ml/kg s.c.) for analgesia. For all experiments, electroencephalography (EEG) was recorded from pairs of stain-less-steel screws (Paul Korth GmbH) placed over the frontal and parietal cortices. To monitor postural tone, nuchal electromyographic (EMG) activity was recorded from three wire electrodes (W3 Wire International) inserted in the dorsal neck musculature. Electrodes were pre-soldered to an ultraminiature pin connector (Preci-Dip) and fixed to the skull with Superbond C&B (Prestige dental) and Paladur dental cement (Kulzer).

### Polysomnographic recordings

After 7-10 days of recover from surgery, single-housed mice were connected to flexible cables allowing free movements. Electrophysiological signals were amplified (Grass Instruments) and analogue-to-digital converted at 512 Hz using SleepSign (Kissei Comtec, Matsumoto, Japan). Habituation was confirmed by verifying 24 h of baseline sleep-wake cycling.

### Offline analysis of polysomnographic data

Polysomnographic recordings were visually scored by 5-s epochs using Sleep Sign software (Vital Recorder, Kissei Comtec). Vigilance states were classified as wake, NREM sleep and REM sleep and cataplexy based on analyses of EEG and EMG recordings. Wake was characterized by a low amplitude, mixed frequency EEG signal in association with a relatively elevated and variable EMG muscle tone and activity. NREM sleep was defined by EEG showing synchronous high amplitude slow wave activity in the delta frequency range (0.5-4 Hz) with a low and stable muscle tone. REM sleep was characterized by theta oscillations (6-9 Hz) and a neck muscle atonia. Power spectral data from 5 s epochs were simultaneously displayed during manual scoring. Transitions to and from NREM or REM sleep were scored based on peak power in the delta or theta bands, respectively. Finally, cataplexy is defined as an abrupt passage from a wake to an EMG showing atonia and an EEG similar to REM sleep or to wake, lasting at least 10s in duration. Cataplexy events were required to be preceded by a wakefulness period lasting at least 40s and are mainly composed of a dominant EEG theta activity as previously described^26^. Moreover, video recordings were used to confirm a matching of EEG and EMG appearance of cataplexy to behavioural immobility and other typical features of cataplexy, like the sudden behavioural arrest, typical of the onset of the event, and the rapid recovery of the muscle tone with behavioral activity at the end of the cataplectic episode.

### Ambient temperature manipulation protocol

A temperature-controlled cabinet was sized to fit 4 plexiglass cages (30 cm x 15 cm) to record 4 mice simultaneously during experiments. To create a Faraday cage, the interior of the cabinet was electrically grounded and lined with reflective aluminum tape. Two infrared (IR) lamps and a convection heat source were used to manipulate the ambient temperature (Ta) of the cabinet under thermostatic control of a programmable timer. The IR lamps were symmetrically positioned to have equal distance from two pairs of cages and they directed toward the ceiling of the cabinet to allow a diffuse reflection of indirect IR light for the rapid warming sessions. The convection heat source was connected to a low pressure forced air circulation system.

Mice were single-housed in individual plexiglass recording cages at constant ambient temperature (23.0 ± 1.0°C) and humidity (40%–60%) under a 12h/12h light/dark cycle (lights on at 7:00 am, or zeitgeber (ZT) time 0). Food and water were available *ad libitum*. After a week of habituation to the cabinet, baseline recordings at constant ambient temperature were performed, followed by a 3–5 day habituation to an Ta manipulation protocol.

The Ta manipulation protocol consists of four bouts of rapid Ta warming performed at two-hour intervals during the middle of either the inactive (light) period (ZT: 2, 4, 6, and 8) or active (dark) period (ZT: 14, 16, 18, 20). Ambient temperatures during the warming bouts ranged between 27.5-32.0°C, producing four 60-min bins in which the Ta reached the high end of the mouse thermoneutral zone (TNZ warm condition) at 32°C. During passive cooling, the four TNZ warm periods were followed by four 60-min bins where the ambient temperatures ranged between 24.5-27.4°C at the low end of the mouse thermoneutral zone (TNZ cool condition). Maximum temperature of 32°C was achieved over the first 30 min during the active warming phase of the warming sessions, followed by passive cooling phases over the next 90 min.

### Food-elicited cataplexy test (FECT)

Hcrt-KO (n=9) mice underwent a food-elicited cataplexy test (FECT) during the dark phase. Each mouse was provided with 3g of milk chocolate as high palatable food in the recording cage immediately prior to lights off (ZT 12). The remaining chocolate was removed from the cage the following day after lights-on (ZT 0). Each mouse underwent the FECT twice at constant ambient temperature and twice during the ambient temperature manipulation protocol, as previously described^27^. The FECT was repeated at least 48 hours apart. Mice were habituated to warm Ta pulsing for 5 days before performing the FECT with Ta manipulation protocol. During FECT, regular food and water were available *ad libitum*.

### Video Actigraphy

Videos of Hcrt-KO mice behaviour were converted to AVI format and uploaded to EthoVision XT (version 10.1.856, Noldus) for movement tracking. On the videos, the area corresponding to the mouse’s cage and nest were manually delimited and calibrated with the software. After setting the optimal detection parameters for a given mouse in each video, the software tracked the animal’s movement from the perspective of the centre point of the animal’s body throughout the entire trial. Each tracked trial was then divided into 1-h time bins, corresponding to the TNZ warming and cooling periods in the analysed video for temperature manipulation conditions. For each time bin, total distance moved, mean and maximum movement velocity parameters were detected and measured, as well as in-zone positioning parameters regarding the nest area: mean time spent within the area with standard deviation and frequency of entering and exiting the nest zone of interest. The software could also generate individual heat maps reflecting time and position within the cage. Data from TNZ warming and cooling conditions were aggregated to obtain a mean value for each parameter. For technical reasons, videos from two of the 9 mice could not be used for the EthoVision tracking system.

### EEG Power Spectral Analyses

Band Power and Spectral power density across behavioral states (Wake, NREM, REM and cataplexy) and experimental conditions (TNZ warm and TNZ cool) were calculated as follows: For each animal and behavioral state, we averaged band power and power spectra across 8 representative TNZ warm bouts and 8 representative TNZ cool bouts per animal. For normalization across animals and subsequent statistical comparisons, band power values in each frequency band (Delta:1-4Hz; Theta:6-9.5Hz; Sigma:10-16Hz; Gamma:30-60Hz) were divided by a normalization factor that corresponds to the sum of the power across all the frequencies according to the experimental condition. The spectral power density values in each bin were then normalized by dividing by the maximal power within the 0.5-20-Hz frequency range in TNZ warm or TNZ cool.

### Statistical Analyses

Statistical analyses were performed comparing the means from each mouse from the three recording conditions, i.e., a control condition at constant temperature (23.0 ± 1.0 °C) and the aggregate of the TNZ warm and TNZ cool conditions. All statistical analyses were performed using GraphPad Prism version 9.2.0 (GraphPad, USA). Two-way repeated measures ANOVA were used for multiple comparisons or a two-tailed paired parametric Student’s t-test for two sample comparisons. Post-hoc ANOVA comparisons were followed by Sidak’s Multiple Comparison Test. Data are presented as the mean ± S.E.M. p values<0.05 were considered to indicate statistical significance.

## Results

To evaluate REM sleep propensity and cataplexy as a function of Ta, mice were exposed to the Ta manipulation protocol involving four bouts of thermoneutral Ta warming occurring at 2-h intervals either during the inactive (light) phase or the active (dark) phase. Continuous electrophysiological recordings allowed for the differentiation of four vigilance states, including Wakefulness, NREM sleep and REM sleep for all mice and the state of cataplexy for Hcrt-KO mice. Figure 2 shows typical continuous sleep-wake recordings from a Hcrt-KO mouse demonstrating either NREM-REM-wake transitions (Fig. 2A) or wake-cataplexy-wake transitions (Fig. 2B). Cataplexy was characterized by bouts of sudden immobility associated with muscle atonia and EEG showing a typical theta activity (Fig. 2B), followed by a rapid transition back to active wakefulness. Cataplexy events were defined according to standard criteria requiring at least 40 seconds of preceding wake and lasting a minimum of 10 seconds in duration^26^.

**Figure 1.**
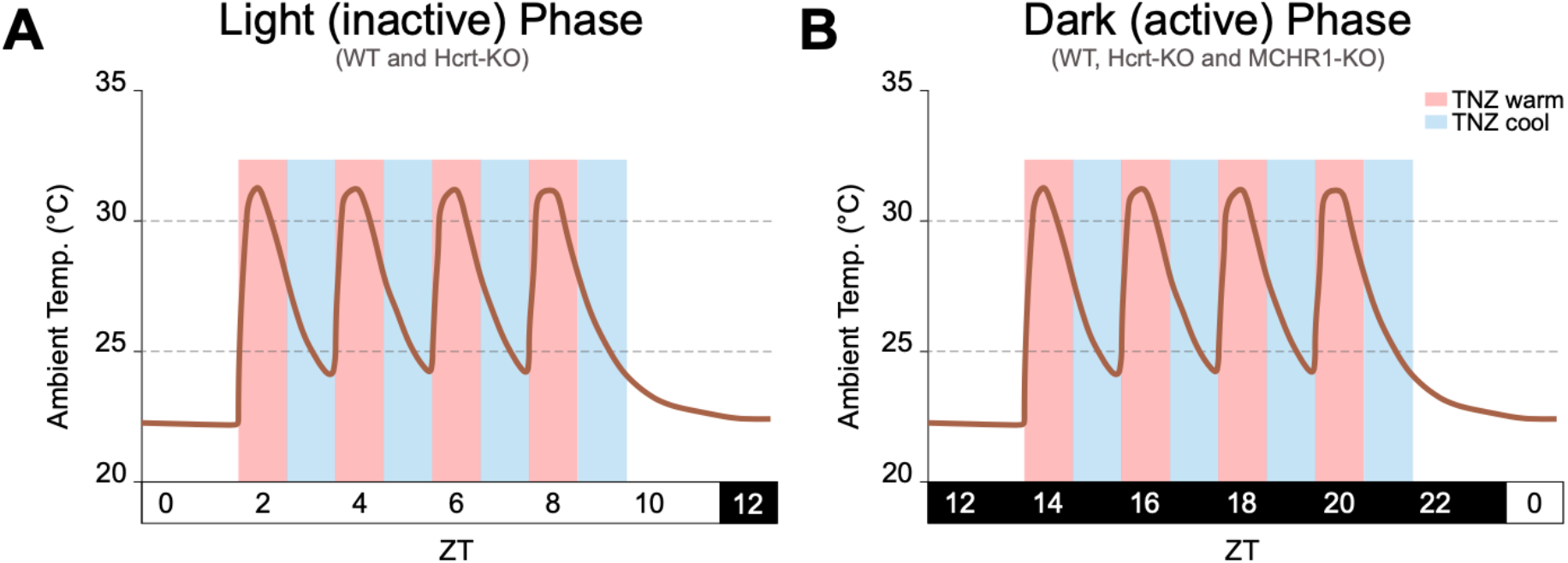
Ambient temperature (Ta) manipulation protocol. Following control recordings at a constant Ta, mice were exposed to four thermoneutral warming pulses presented at 2-h intervals during either **A)** the middle of the inactive (light) phase starting at zeitgeber time (ZT) 2 or **B)** in the middle of the active (dark) phase beginning at ZT 14. This protocol provided alternating 60-min bouts of thermoneutral zone (TNZ) warm (red vertical bars; Ta approx. 27.5°-32.0°C) and TNZ cool (blue; Ta approx. 24.5°C-27.5°C) conditions. Wt and Hcrt-KO mice were exposed to both circadian Ta manipulation conditions, whereas the MCHR1-KO mice were only exposed to Ta manipulation during the dark phase. All mice were also recorded in constant 23.0 ± 1.0°C baseline conditions (not shown).

**Figure 2.**
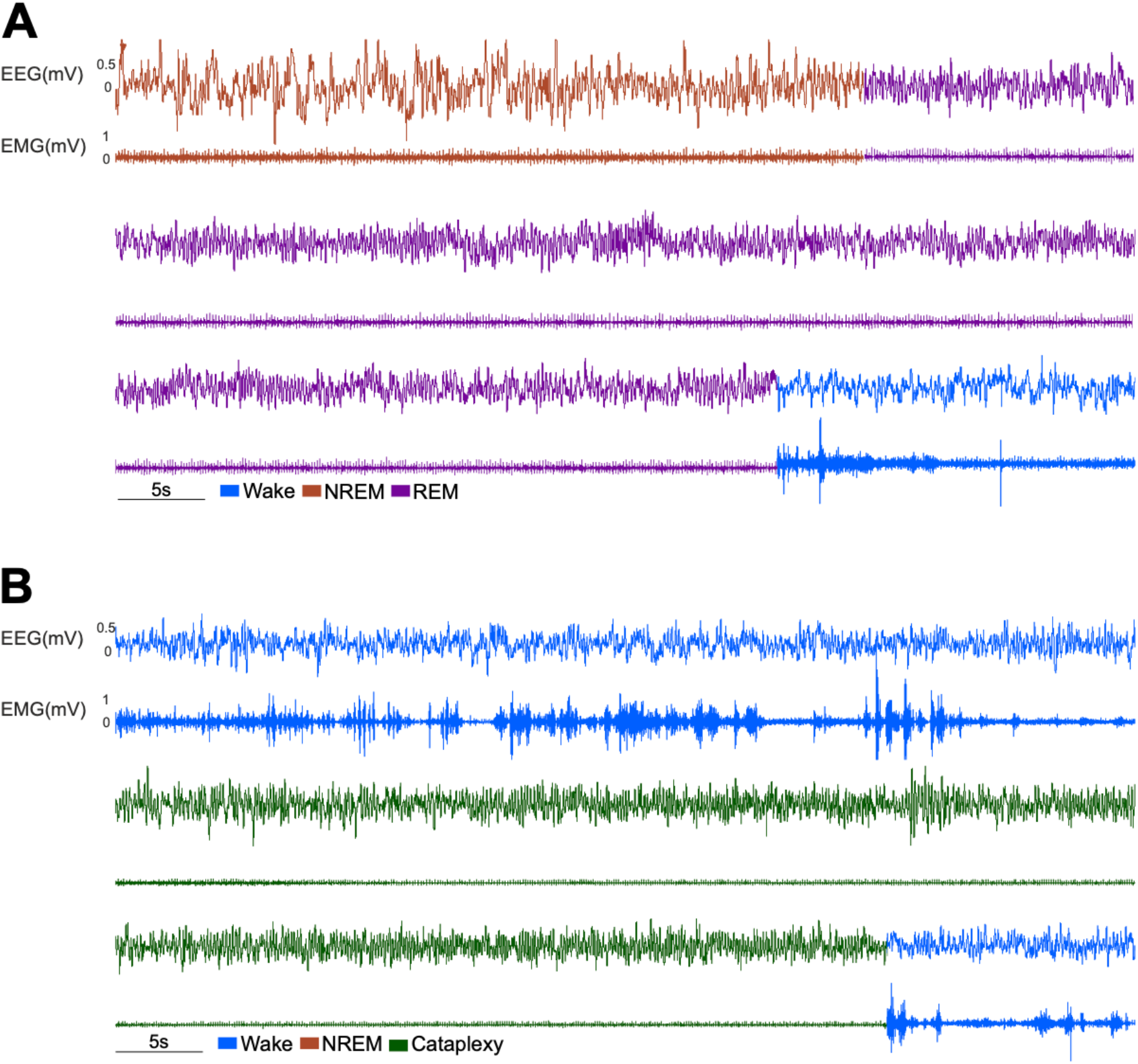
Physiological electroencephalographic (EEG) and electromyographic (EMG) recordings from a hypocretin knockout (Hcrt-KO) mouse. **A)** A typical continuous electrophysiological recording showing NREM (red)-REM (purple)-Wake (blue) transitions with high amplitude slow-waves and reduced EMG tone during NREM sleep, EEG theta rhythm with muscle atonia during REM sleep and sudden increase in EMG activity with mixed frequency EEG activity during Wake. **B)** A cataplexy event (green) shows similar EEG and EMG activity as seen during REM sleep but transitioning abruptly from Wake.

### REM Sleep Propensity and Sleep-Wake expression

Opposite responsiveness patterns with respect to wakefulness, NREM sleep and REM sleep were observed between WT and Hcrt-KO mice as a function of both circadian time and Ta manipulation (Fig. 3). With respect to REM sleep propensity, we found that thermoneutral Ta warming increased REM sleep expression in WT mice selectively during the light phase with no effect when presented during the dark phase (see Fig. 3C vs 3F), as was previously demonstrated^25^. Hcrt-KO mice, however, showed an opposite circadian responsiveness pattern to Ta manipulation. As shown in figure 3C, REM sleep expression in Hcrt-KO mice significantly decreased during Ta warming presented during the light phase compared to the baseline condition. Moreover, Hcrt-KO mice during the dark circadian phase showed a trend to increase total REM sleep during Ta warming compared to WT mice (Fig. 3F).

**Figure 3.**
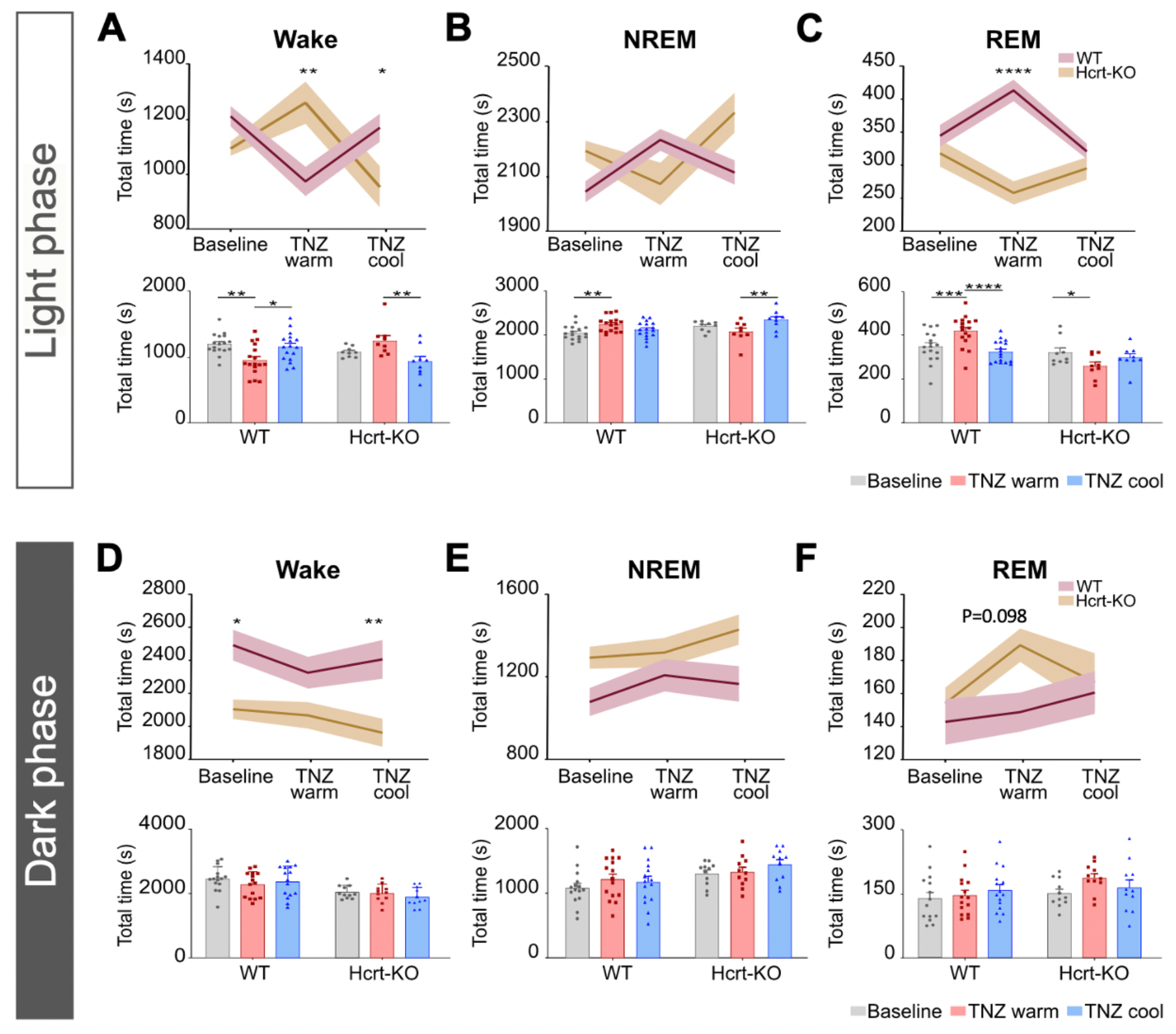
Sleep-wake expression as a function of ambient temperature (Ta) and circadian time. **A-C)** Wake, NREM sleep and REM sleep expression during the inactive (light) phase in wild type (WT, n=17) and hypocretin-Knockout (Hcrt-KO, n=9) mice across baseline (23.0 ± 1.0°C), thermoneutral zone (TNZ) warm and TNZ cool conditions. The upper graphs show between group (WT vs Hcrt-KO) comparisons, whereas the lower graphs show within group comparisons. **D-F)** Using the same methodology, total Wake, NREM sleep and REM sleep expression is shown for the active (dark) circadian phase between WT (n=15) and Hcrt-KO (n=11) mice. Data were analyzed using two-way ANOVA and post-hoc Sidak’s comparison test. Data are presented as means ± standard error of the mean (SEM). *p<0.05; **p<0.01; ***p<0.001; ****p<0.0001.

When comparing TNZ warm vs TNZ cool conditions during the light circadian phase, WT mice showed an increased number of REM sleep bouts during the Ta warm condition as previously demonstrated. However, mean REM sleep bout durations remained unchanged as a function of Ta in both groups of mice (Suppl. Fig. 1). Finally, our prior results demonstrated that MCHR1-KO mice fail to show changes in REM sleep expression during Ta warming during the light phase^25^. We performed the same Ta manipulation protocol in MCHR1-KO mice during the dark phase and again found no changes to REM sleep as a function of Ta (Suppl. Fig. 2).

Opposite circadian responsiveness patterns between WT and Hcrt-KO mice were also observed for Wake and NREM sleep. Specifically, WT mice during the light phase showed a significant decrease in total wake time during thermoneutral Ta warming compared to both the baseline and TNZ cool conditions (Fig. 3A). Hcrt-KO mice, in contrast, showed a significant increase in total wake time during Ta warming compared to TNZ cool condition (Fig. 3A) manifested primarily by an increase in wake bout duration without a significant change in wake bout number (Suppl. Fig. 1). NREM sleep appeared to be less affected by Ta condition in all groups of mice. For example, although WT mice increased NREM sleep during Ta warming with respect to the baseline condition during the light phase, a post-hoc analysis showed no differences with respect to the TNZ cool condition (Fig. 3B). In contrast, Hcrt-KO mice again showed an opposite pattern with a significant decrease in NREM sleep durations during Ta warming compared to the cool condition (Fig. 3B). During the active (dark) phase, trends for opposite responsiveness patterns were again observed for wake and NREM sleep in the WT and Hcrt-KO groups (Fig. 3D and 3E) but did not reach significance. Finally, NREM sleep bout number and mean NREM bout durations were not significantly changed between the TNZ warm and cool conditions in all groups of mice during either the light or dark phases (Suppl. Fig. 1).

Taken together, these data show that responsiveness to Ta warming is dependent on circadian time and shows an opposite pattern between WT and Hcrt-KO mice. Moreover, mice lacking the MCHR1 receptor, in contrast, fail to show changes in REM sleep during either circadian phase, consistent with the critical role of the MCH system in driving increased REM sleep expression during thermoneutral Ta warming as was previously demonstrated^25^.

### Cataplexy expression and Ta manipulation

To examine the role of Ta manipulation on both spontaneous and food-elicited cataplexy, the Ta manipulation protocol was applied during the active (dark) circadian phase. Spontaneous cataplexy was evaluated using the protocol described above, whereas the food-elicited cataplexy test (FECT) was performed on a separate night using a standard protocol of placing 3g of milk chocolate in the cage at ZT 12 immediately prior to lights off, corresponding two hours prior to the start of the Ta manipulation protocol. The remainder of milk chocolate was removed in the morning upon lights on.

As expected, the presence of chocolate markedly increased the total number of cataplexy events in Hcrt-KO mice over the normal chow condition (see Fig. 4A and 4B). Moreover, Ta warming significantly and consistently decreased cataplexy expression compared to baseline (Fig. 4A) and the TNZ cool conditions (Fig. 4B). Although the effect of thermoneutral Ta warming on decreasing cataplexy was most pronounced in the FECT condition, similar results were observed for spontaneous cataplexy (Fig. 4B). The reduction in cataplexy during Ta warming in the FECT condition was primarily driven by a decrease in the total number of cataplexy bouts (Fig. 4B). However, Ta cooling in the FECT condition was associated with a significant increase in mean cataplexy bout duration compared to baseline (Fig. 4B).

**Figure 4.**
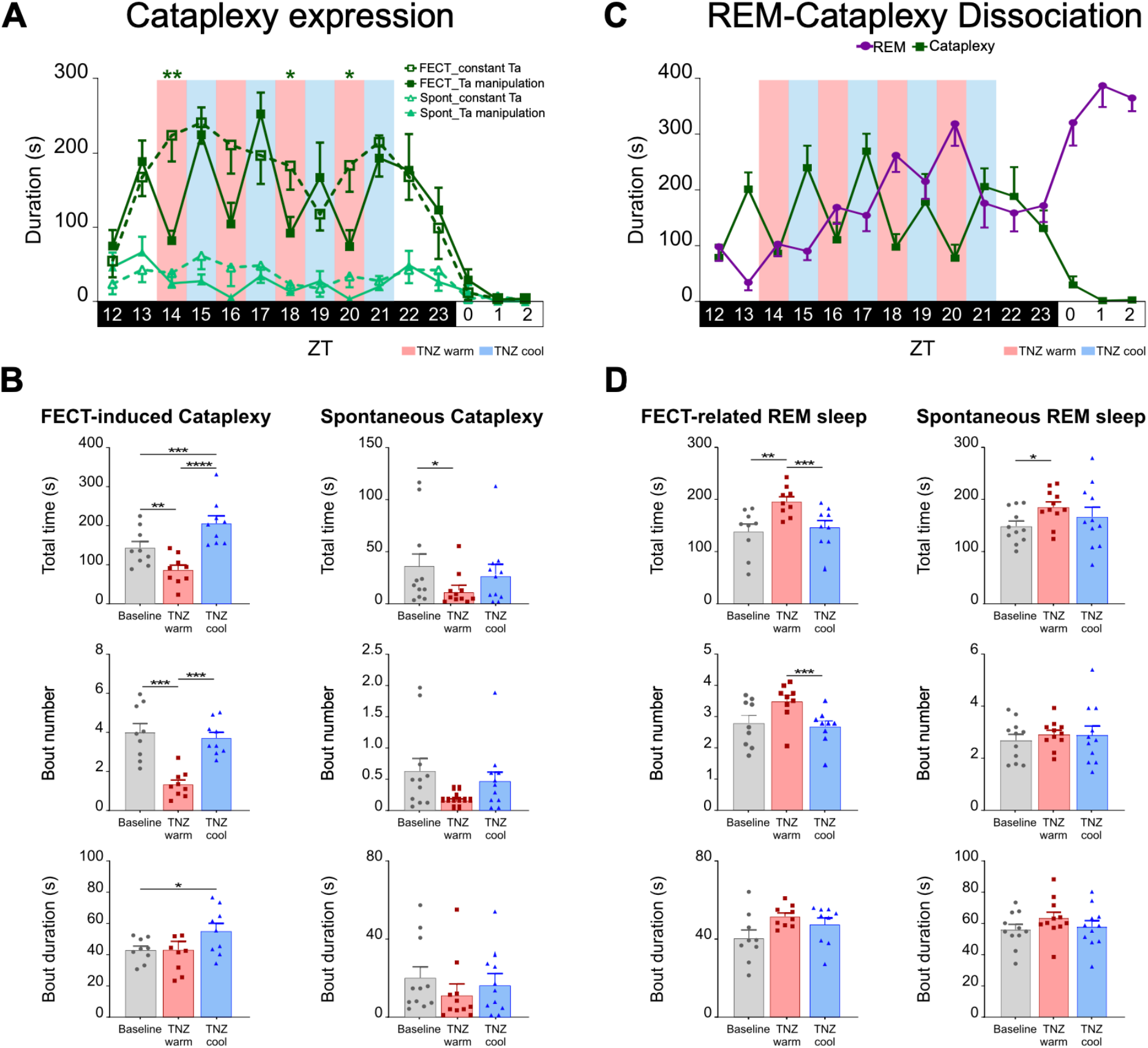
Dissociation of cataplexy and REM sleep during thermoneutral ambient temperature (Ta) warming in Hcrt-KO (n=9) mice. **A)** Cataplexy expression in 1-h bins during the 12-h dark phase, followed by the first 3-h of the light phase showing results during the food elicited cataplexy test (FECT, dark green) and spontaneous (light green) cataplexy conditions either during constant Ta (dashed line) or the Ta manipulation protocol (solid line). Significant differences for the FECT condition using a two-tailed student’s t test between the constant Ta and Ta manipulation conditions are shown. **B)** Total durations of cataplexy, cataplexy bout number and mean cataplexy bout durations per hour for the baseline, TNZ warm and TNZ cool conditions. The FECT and spontaneous cataplexy recordings were performed on separate nights. **C)** The dissociation of REM sleep and cataplexy is demonstrated as a temperature effect during the dark phase with Ta manipulation and as a circadian effect during the light phase. REM sleep expression during the FECT condition is displayed with a replotting of cataplexy expression from A) across the 12-h dark phase and the first 3-h of the light phase. **D)** Total durations of REM sleep, REM sleep bout number and mean REM sleep bout durations expressed per hour of baseline, TNZ warm and TNZ cool conditions. Results were obtained on separate nights for the presence of chocolate (FECT) or without chocolate (spontaneous) conditions. Data were analyzed using two-way ANOVA and post-hoc Sidak’s comparison test. Data are presented as means ± SEM. *p<0.05; **p<0.01; ***p<0.001; ****p<0.0001.

Ta manipulation coupled with FECT also clearly demonstrated significant changes with respect to REM sleep expression, showing an opposite responsiveness pattern from cataplexy. As figure 4C shows, Ta warming significantly increased REM sleep expression compared to both the TNZ cool and baseline conditions. This effect was more pronounced in the second half of the night where REM sleep expression is particularly increased in Hcrt-KO mice (Fig. 4C).

These data demonstrate that thermoneutral Ta warming triggered a marked dissociation between cataplexy and REM sleep where total REM sleep time significantly increased concomitant with cataplexy reduction. The increase in REM sleep was primarily driven by a significant increase in the total number of REM sleep bouts. Finally, in addition to a Ta manipulation effect during the dark phase, the dissociation of REM sleep and cataplexy was also observed as a circadian effect during the light phase at constant Ta in both the baseline and FECT conditions (Fig. 4C).

### Video actigraphy and sleep-wake behavior

Next, we determined whether the decrease in cataplexy and increase in REM sleep observed during Ta warming could be explained by a change in waking behavior or sleep-wake expression. Therefore, EEG/EMG recordings and detailed video actigraphy analyses of narcoleptic Hcrt-KO mice were performed to directly compare the TNZ warm and TNZ cool conditions. We found no significant changes in total wake durations during the active (dark) phase in Hcrt-KO mice as a function of Ta condition (see Fig. 3D and 3E). Similarly, no significant differences were observed between the warm vs cool conditions for the total number of wake bouts mean wake bout durations (Suppl. Fig. 1). Moreover, there were no significant differences observed for total NREM sleep time, number of NREM sleep bouts, or mean NREM sleep bout durations as a function of Ta manipulation during the dark phase (Fig. 3E and Suppl. Fig. 1).

Video recordings were then analyzed in detail using Ethovision tracking software. Individual heat maps reflecting time and position in the cage failed to identify qualitative differences between the TNZ warm vs TNZ cool conditions (see Fig. 5A vs 5B). Moreover, no differences were observed between Ta conditions for any quantitative variable examined, including distance moved, mean velocity, time in nest, or frequency of entering the nest (Fig. 5C-5F). Taken together, these data indicate that the decrease in cataplexy and increase in REM sleep expression as a function of Ta was independent of both waking and NREM sleep and could not be explained by changes in waking motor activity.

**Figure 5.**
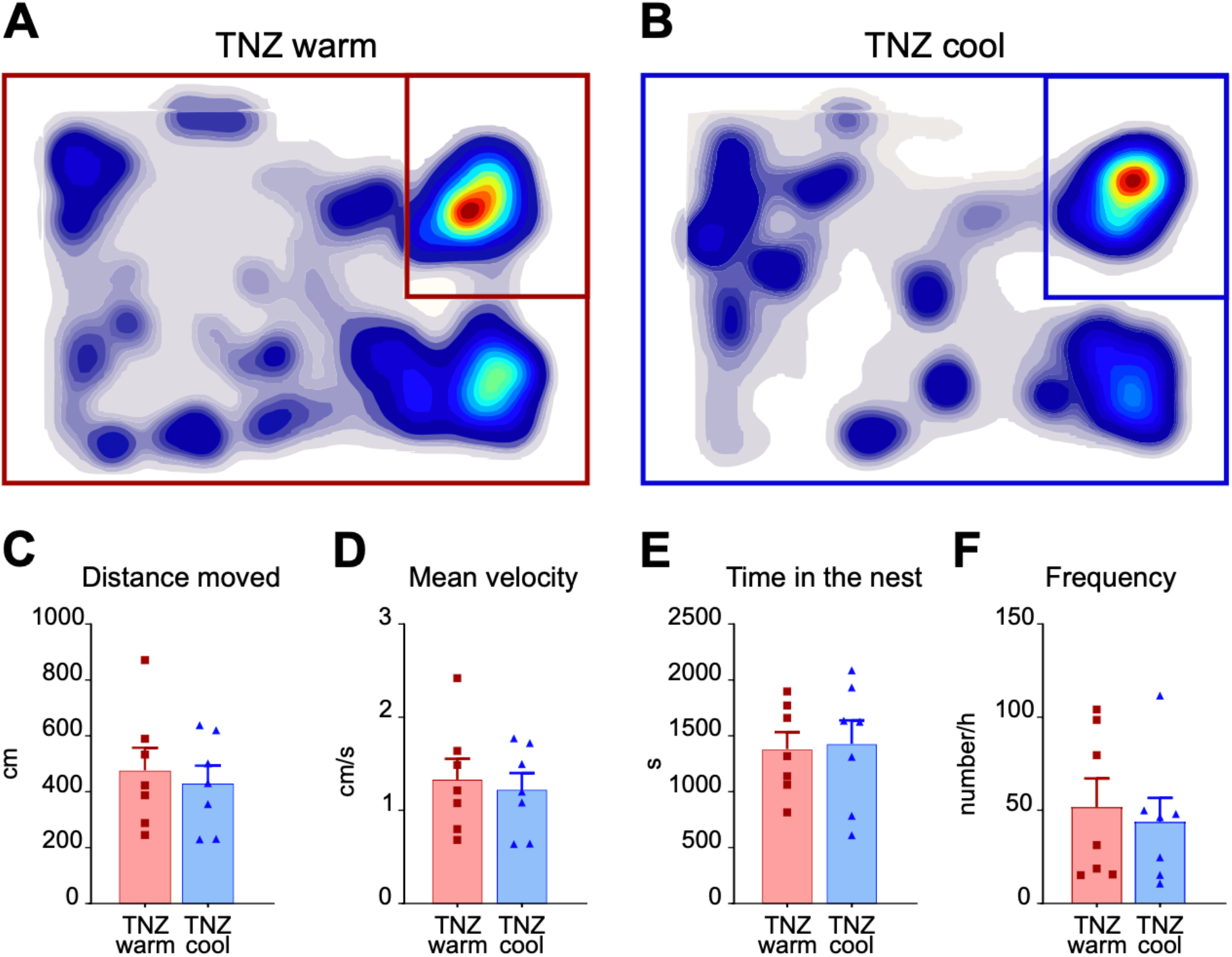
Video actigraphy of Hcrt-KO (n=7) mice during the food elicited cataplexy test (FECT) comparing the TNZ warm and TNZ cool conditions. **A-B)** Individual heat maps failed to identify qualitative differences with respect to time and position in the cage comparing the TNZ warm vs TNZ cool conditions as shown here for a representative Hcrt-KO mouse. The box in the upper right corner represents the position of the nest. **C-F)** Quantitative analyses regarding distance moved, mean velocity, time in nest and frequency of entering the nest are shown. No significant differences were observed between the TNZ warm and TNZ cool conditions.

### Power Spectral EEG Analyses

Power spectral density and band power were analyzed for both WT and Hcrt-KO mice during the active (dark) phase comparing the TNZ warm and TNZ cool conditions. For wake and NREM sleep, no differences were observed as a function of Ta for either group of mice with the exception of an isolated and small increase at the lowest frequencies in delta power (1-4 Hz) for the WT group during the TNZ cool condition (Suppl. Fig. 3).

REM sleep revealed several differences with respect to band power and power spectral analyses as a function of Ta condition. Specifically, WT mice showed a small but significant increase in the delta power band in REM sleep during the Ta warm condition (Fig. 6A) which was not observed in Hcrt-KO mice (Fig. 6B). Although no significant differences were observed for the theta band analysis (6-9.5 Hz band), small but significant increases in spectral power were observed for individual frequencies in the theta frequency range during REM sleep in the TNZ warm condition for both the WT and Hcrt-KO groups (Fig. 6A and 6B).

**Figure 6.**
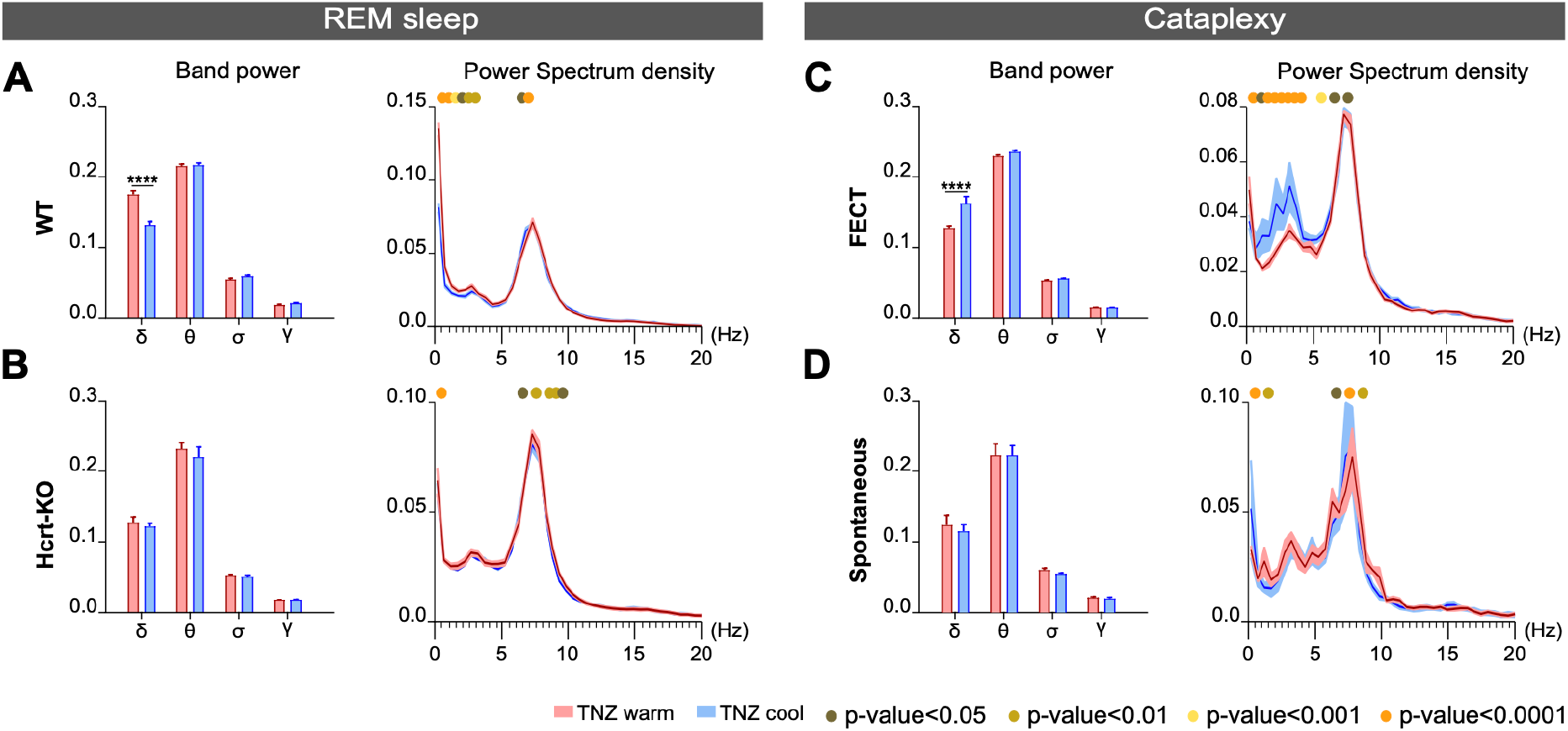
Relative EEG power during REM sleep and cataplexy in the TNZ warm (red) vs TNZ cool (blue) conditions. **A-B)** Band power and power spectral analyses for REM sleep in WT (n=15) and Hcrt-KO mice (n=11). **C-D)** Band power and spectral power density during cataplexy in either the food elicited cataplexy test (FECT, n=8) in the presence of highly palatable chocolate (A) or during spontaneous cataplexy (n=11) in the absence of chocolate (B). Levels of significance indicated by color coding in the figure.

The most striking difference between the TNZ warm and TNZ cool conditions was observed during food-elicited cataplexy for the delta band during the Ta cool condition in Hcrt-KO mice (Fig. 6C). This increase in delta power was consistently observed across delta frequencies. These results show similarities to the “delta-theta” sleep events described in dual Hcrt-MCH ablated mice as seen during sudden episodes of immobility^16^. Similar findings, however, were not observed in the delta band for the spontaneous cataplexy condition (Fig. 6D).

## Discussion

It is well established that neural mechanisms controlling both sleep and thermoregulation are tightly integrated^28–32^. For example, across mammalian species examined, thermoneutral Ta warming is known to preferentially increase REM sleep over NREM sleep^25,33–35^. Moreover, REM sleep in endotherms is characterized by a loss of thermoregulatory defenses such as panting, shivering and sweating^30,36^. Our recent work demonstrates a critical role of the MCH system in dynamically increasing REM sleep expression when the need for thermoregulatory defense is minimized such as when mice are exposed to brief thermoneutral Ta warming bouts^25^. This prior work confirms the presence of neural mechanisms that favor REM sleep when the need for thermoregulatory defense is reduced or absent.

Here, we demonstrate that narcoleptic mice show abnormal responsiveness patterns regarding REM sleep and wakefulness expression in response to Ta warming compared to WT mice. Whereas WT mice increased REM sleep and decreased wakefulness during Ta warming presented in the inactive (light) phase, Hcrt-KO mice show an opposite circadian pattern with decreased REM sleep and increased wakefulness. Interestingly, Hcrt-KO mice showed increased REM sleep in response to Ta warming presented in the dark phase, although WT mice do not alter REM sleep or wake expression in the same condition. Finally, we observed a markedly decreased cataplexy expression during thermoneutral Ta warming concomitant with a significant increase in REM sleep even though waking activity remained unchanged. This ability to dissociate REM sleep and cataplexy using Ta manipulation was previously unknown, a finding that we hypothesize may present a new technique to dissect neural circuits responsible for the expression of these two phenomena.

In our study we limited Ta manipulation to the narrow thermoneutral zone (TNZ) of mice, i.e., between 24.5-32°C. It remains to be determined how Ta manipulation either below or exceeding this narrow range may impact either REM sleep or cataplexy. Prior work comparing the housing of narcoleptic mice at a constant low of 20°C vs a high at 30°C shows that both NREM and REM sleep are increased during the warm vs cool condition^37^. However, a Ta of 20°C is well below the thermoneutral zone for mice, thus making comparisons with our data difficult. For our experiments, we specifically restricted Ta manipulation to within the TNZ so as to minimize any confounding effects from NREM sleep expression. As a result, our protocol resulted in minimal or small changes to NREM sleep between the TNZ warm and TNZ cool conditions, demonstrating that the alterations in REM sleep expression were relatively independent of other sleep-wake parameters.

### Link between narcolepsy and abnormal thermoregulation

The Hcrt system is known to play an important role in not only driving wakefulness, but also in increasing core body temperature associated with the waking state. Hcrt neurons, for example, increase brown adipose tissue (BAT) activity through their direct projection to the raphe pallidus^38,39^. The Hcrt system is also known to activate the sympathetic nervous system^40–42^ and may thus also play a role in peripheral vasoconstriction during wakefulness so as to decrease heat loss and improving heat retention.

Prior work has also shown that patients with narcolepsy show abnormalities in thermoregulatory responses compared to healthy controls. For example, healthy individuals usually show cool distal skin temperatures during active wakefulness associated with the circadian day, but then show vasodilatation and distal extremity warming in the evening both in anticipation of sleep and during sleep. In contrast, patients with narcolepsy demonstrate an opposite thermoregulatory pattern with distal skin warming during active daytime that is normally associated with sleep and correlating with severity of daytime sleepiness^43,44^. Moreover, whereas daytime distal skin cooling in patients with narcolepsy significantly improves their ability to maintain wakefulness, proximal skin warming at night improves their sleep consolidation and slow-wave sleep expression^45,46^. Interestingly, distal skin warming presented at night in human narcolepsy appears to paradoxically increase wakefulness^45^, contrary to the increased sleep propensity from skin warming observed in healthy controls^47–49^. Consistent with this prior work, our narcoleptic mice also showed a paradoxical increase in waking when Ta warming was presented during their inactive phase, in contrast to decreased waking in WT mice.

Our findings of opposite responsiveness patterns in mice with respect to REM sleep and wakefulness expression further support a dysregulation of thermoregulatory responses in narcolepsy. We hypothesize that the lateral hypothalamus plays a key role in integrating numerous physiological variables, including energy status, circadian time, sleep homeostasis and ambient temperature for the output control of wakefulness vs REM sleep^50^. The mechanism by which the loss of the Hcrt neuropeptide leads to these thermoregulatory changes requires further investigation. However, we hypothesize that in addition to disrupting output control mechanisms, Hcrt loss likely compromises complex LH circuits required for the integration of these key input physiological variables.

### REM sleep propensity and narcolepsy

Given the known reciprocal activity patterns of Hcrt and MCH neurons suggesting either direct or indirect reciprocal inhibition^19^, we originally hypothesized that Hcrt loss would lead to increased REM sleep propensity secondary to MCH disinhibition. Specifically, we hypothesized that Hcrt-KO mice would show even greater increases in REM sleep in response to thermoneutral Ta warming secondary to a hyperactive responsiveness in the MCH system. Indeed, this concept of disinhibition of the MCH system in narcolepsy was previously suggested using chemogenetic manipulation of MCH neurons in Hcrt-KO mice^14^. However, our results suggest a more complicated relationship between REM sleep expression and Ta condition.

On the one hand, Hcrt-KO mice show increased REM sleep propensity compared to WT mice when thermoneutral Ta warming is specifically presented during the active (dark) phase. On the other hand, Hcrt-KO mice not only fail to increase REM sleep during Ta warming presented during the inactive (light) phase, but unexpectedly show a circadian specific decrease in REM sleep in response to Ta warming with increased wakefulness. Although currently speculation, one possible mechanism may be that MCH neurons are already near their maximum firing responsiveness level during the inactive phase in the absence of Hcrt. Thus, they may be unable to mount further increased responsiveness to small increases in thermoneutral Ta warming. In the active dark phase, however, we speculate that the MCH system may have a lower baseline responsiveness level, allowing Hcrt-KO mice to mount a response to Ta warming that is normally prevented or counter-balanced during the circadian active phase in the presence of Hcrt in WT mice.

The use of Ta warming to selectively increase REM sleep expression during the circadian active phase in narcoleptic mice presents a potential novel translational application for human narcolepsy that requires further investigation. The characteristic increase in REM sleep propensity is a key parameter used as part of the multiple sleep latency test (MSLT) to diagnose narcolepsy^3,51^, but this clinical test has been criticized as lacking sufficient sensitivity and specificity^52–54^. Even for patients with unequivocal narcolepsy with cataplexy, a significant subpopulation fail to show the minimum requirement of at least two naps with REM sleep^55^. Moreover, test-retest reliability of REM sleep expression for patients initially diagnosed with either narcolepsy without cataplexy or idiopathic hypersomnia is poor with approximately 50% of patients showing a change in these diagnoses following a retesting with the MSLT^3,52,53^. These data confirm a clinical need for more robust diagnostic tools in supporting the diagnosis of narcolepsy, particularly narcolepsy without cataplexy, and in differentiating such patients from other hypersomnias^56^. We hypothesize that thermoneutral ambient or skin temperature warming during MSLT testing could present a novel technique to increase both sensitivity and specificity of the test in the diagnosis of narcolepsy, a hypothesis requiring further investigation.

### Ambient temperature and cataplexy

Thermoneutral Ta manipulation presents an unexpected ability to dissociate REM sleep and cataplexy where Ta warming increases REM sleep but decreases cataplexy. Although novel, a similar dissociation between REM sleep and cataplexy can be observed in Hcrt-KO mice during the circadian inactive (light) phase in constant Ta conditions. Indeed, cataplexy expression during the circadian inactive phase is rarely observed in narcoleptic mice. Although these data may seem to support recent findings suggesting that REM sleep and cataplexy may be independent behavioral states with unique neural mechanisms^6^, this perspective with respect to our findings requires further investigation.

The behavioral state instability hypothesis suggests that REM sleep components such as muscle atonia intrude into the waking state. We speculate that REM sleep and cataplexy may be dissociated in narcolepsy through a gating of MCH neuronal activity. Specifically, our data are consistent with the hypothesis that cataplexy is favored to occur in cooling when both Hcrt and MCH activities are likely to be minimized in Hcrt-KO mice during Ta cooling. The normal response to Ta cooling is arousal/waking, a behavioral response normally associated with increased Hcrt activity and a decrease in MCH activity^19^. Ta cooling, therefore, may lead to a dissociated condition not normally seen in WT mice, a condition resulting from the concomitant hypoactivity of both the MCH and Hcrt systems (Fig. 7). In support of this hypothesis, it has recently been demonstrated that cataplexy expression is markedly increased in Hcrt and MCH double-ablated mice^16^. Moreover, calcium imaging using the Inscopix miniscope shows that cataplexy is associated with a general absence of MCH activity^15^. These data suggest that inactivity of both the Hcrt and MCH systems may further exacerbate behavioral state instability and thus favor cataplexy.

**Figure 7.**
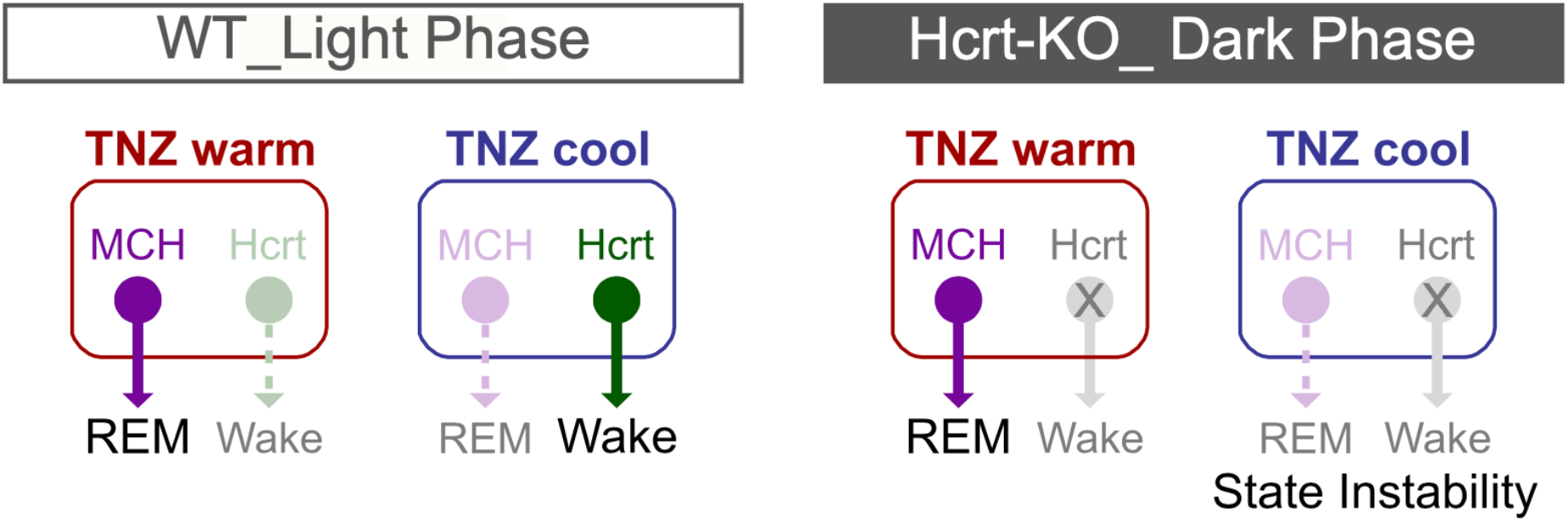
Theoretical construct on the role of the MCH and Hcrt systems in modulating dynamic state stability as a function of Ta. For WT mice, increased REM sleep expression during Ta warming is driven by MCH neuronal activity25, whereas the Hcrt system may favor wakefulness during Ta cooling. In Hcrt-KO mice, Ta warming may theoretically favor REM sleep expression through an MCH response, thus maintaining state stability. Ta cooling, however, may lead to state instability if the activity of MCH system is decreased without the counterbalance of increased Hcrt activity.

Recent work further shows a peculiar state of motor immobility in dual MCH-Hcrt ablated mice associated with marked increase in delta power reported as “delta-theta” sleep^16^. Interestingly, we observed a significant increase in delta power in Hcrt-KO mice during cataplexy specifically associated with Ta cooling. To what extent this increase in delta activity may reflect hypoactivity of the MCH system during Ta cooling remains to be determined.

### Summary

Our findings demonstrate that Hcrt-KO mice show opposite circadian responsiveness patterns compared to WT mice with respect to REM sleep and wake expression as a function of Ta. Given that patients with narcolepsy also exhibit altered thermoregulatory response patterns, we suggest that translational approaches utilizing Ta manipulation to improve sensitivity and specificity REM sleep expression in diagnostic testing for narcolepsy merits further investigation. Moreover, the ability to dissociate REM sleep and cataplexy in Hcrt-KO mice as a function of Ta may provide new avenues to dissect neural circuits controlling these two phenomena. Indeed, further work is required to determine the role, if any, of the MCH system in gating cataplexy expression.

## Acknowledgements

We would like to thank Noemi Komagata, Blerina Latifi and Lucia Di Stefano for their specific contributions toward either data acquisition or data analyses. We would also like to thank members of the ZEN laboratory for their technical assistance, including Thomas Rusterholz for MatLab code and help with computational analyses. This work is supported by the Center for Experimental Neurology and the Department of Neurology at the University of Bern, Bern University Hospital, Inselspital, as well as the Interfaculty Research Grant (IRC), the Sleep Medicine Research Foundation and the Ohio Sleep Medicine Institute. AA was supported by the Human Frontier Science Program (RGY0076/2012), Inselspital University Hospital Bern, Swiss National Science Foundation (31003A_156156), European Research Council (725850), Sinergia (CRSII3_160803), and the University of Bern.

## Author Contributions

BV, LB and MHS performed experiments, analyzed data, performed statistical analyses and generated figures. SB performed the EthoVision video actigraphy analyses. MHS designed the experiments. MHS and BV wrote the manuscript. All authors contributed to manuscript revisions. MHS conceived the project and directly supervised the work.

## References

1. Chemelli, R.M., Willie, J.T., Sinton, C.M., Elmquist, J.K., Scammell, T., Lee, C., Richardson, J.A., Clay Williams, S., Xiong, Y., Kisanuki, Y., et al. (1999). Narcolepsy in orexin knockout mice: Molecular genetics of sleep regulation. Cell 98, 437–451.

2. Peyron, C., Faraco, J., Rogers, W., Ripley, B., Overeem, S., Charnay, Y., Nevsimalova, S., Aldrich, M., Reynolds, D., Albin, R., et al. (2000). A mutation in a case of early onset narcolepsy and a generalized absence of hypocretin peptides in human narcoleptic brains. Nat. Med. 6, 991–997.

3. Bassetti, C.L.A., Adamantidis, A., Burdakov, D., Han, F., Gay, S., Kallweit, U., Khatami, R., Koning, F., Kornum, B.R., Lammers, G.J., et al. (2019). Narcolepsy — clinical spectrum, aetiopathophysiology, diagnosis and treatment. Nat Rev Neurol 15, 519–539.

4. Mahoney, C.E., Cogswell, A., Koralnik, I.J., and Scammell, T.E. (2019). The neurobiological basis of narcolepsy. Nat Rev Neurosci 20, 83–93.

5. Schoch, S.F., Werth, E., Poryazova, R., Scammell, T.E., Baumann, C.R., and Imbach, L.L. (2017). Dysregulation of Sleep Behavioral States in Narcolepsy. Sleep 40.

6. Vassalli, A., Dellepiane, J.M., Emmenegger, Y., Jimenez, S., Vandi, S., Plazzi, G., Franken, P., and Tafti, M. (2013). Electroencephalogram paroxysmal theta characterizes cataplexy in mice and children. Brain 136, 1592–1608.

7. Mochizuki, T. (2004). Behavioral State Instability in Orexin Knock-Out Mice. Journal of Neuroscience 24, 6291–6300.

8. Vu, M.H., Hurni, C., Mathis, J., Roth, C., and Bassetti, C.L. (2011). Selective REM sleep deprivation in narcolepsy. J Sleep Res 20, 50–56.

9. Okura, M. (2000). Sulpiride, a D2/D3 Blocker, Reduces Cataplexy but not REM Sleep in Canine Narcolepsy. Neuropsychopharmacology 23, 528–538.

10. Kantor, S., Mochizuki, T., Janisiewicz, A.M., Clark, E., Nishino, S., and Scammell, T.E. (2009). Orexin neurons are necessary for the circadian control of REM sleep. Sleep 32, 1127–1134.

11. Selbach, O., and Haas, H.L. (2006). Hypocretins: The timing of sleep and waking. Chronobiology International 23, 63–70.

12. Roman, A., Meftah, S., Arthaud, S., Luppi, P.-H., and Peyron, C. (2018). The inappropriate occurrence of rapid eye movement sleep in narcolepsy is not due to a defect in homeostatic regulation of rapid eye movement sleep. Sleep 41.

13. Willie, J.T., Takahira, H., Shibahara, M., Hara, J., Nomiyama, M., Yanagisawa, M., and Sakurai, T. (2011). Ectopic Overexpression of Orexin Alters Sleep/Wakefulness States and Muscle Tone Regulation during REM Sleep in Mice. J Mol Neurosci 43, 155–161.

14. Naganuma, F., Bandaru, S.S., Absi, G., Mahoney, C.E., Scammell, T.E., and Vetrivelan, R. (2018). Melanin-concentrating hormone neurons contribute to dysregulation of rapid eye movement sleep in narcolepsy. Neurobiol. Dis. 120, 12–20.

15. Sun, Y., and Liu, M. (2020). Hypothalamic MCH Neuron Activity Dynamics during Cataplexy of Narcolepsy. eNeuro 7, ENEURO.0017-20.2020.

16. Jung Hung, C., Ono, D., Kilduff, T., and Yamanaka, A. (2019). Dual Orexin and MCH neuron-ablated mice display severe sleep attacks and cataplexy. Journal of the Japan Welding Society 88, 427–434.

17. Ferreira, J.G.P., Bittencourt, J.C., and Adamantidis, A. (2017). Melanin-concentrating hormone and sleep. Current Opinion in Neurobiology 44, 152–158.

18. Peyron, C., Tighe, D.K., van den Pol, A.N., de Lecea, L., Heller, H.C., Sutcliffe, J.G., and Kilduff, T.S. (1998). Neurons Containing Hypocretin (Orexin) Project to Multiple Neuronal Systems. The Journal of Neuroscience 18, 9996–10015.

19. Hassani, O.K., Lee, M.G., and Jones, B.E. (2009). Melanin-concentrating hormone neurons discharge in a reciprocal manner to orexin neurons across the sleep-wake cycle. Proc. Natl. Acad. Sci. U.S.A. 106, 2418–2422.

20. Jego, S., Glasgow, S.D., Herrera, C.G., Ekstrand, M., Reed, S.J., Boyce, R., Friedman, J., Burdakov, D., and Adamantidis, A.R. (2013). Optogenetic identification of a rapid eye movement sleep modulatory circuit in the hypothalamus. Nature Neuroscience 16, 1637–1643.

21. Vetrivelan, R., Kong, D., Ferrari, L.L., Arrigoni, E., Madara, J.C., Bandaru, S.S., Lowell, B.B., Lu, J., and Saper, C.B. (2016). Melanin-concentrating hormone neurons specifically promote rapid eye movement sleep in mice. Neuroscience 336, 102–113.

22. Li, S.-B., Nevárez, N., Giardino, W.J., and de Lecea, L. (2018). Optical probing of orexin/hypocretin receptor antagonists. Sleep 41.

23. Adamantidis, A.R., Schmidt, M.H., Carter, M.E., Burdakov, D., Peyron, C., and Scammell, T.E. (2020). A circuit perspective on narcolepsy. Sleep 43, zsz296.

24. Adamantidis, A.R., Zhang, F., Aravanis, A.M., Deisseroth, K., and de Lecea, L. (2007). Neural substrates of awakening probed with optogenetic control of hypocretin neurons. Nature 450, 420–424.

25. Komagata, N., Latifi, B., Rusterholz, T., Bassetti, C.L.A., Adamantidis, A., and Schmidt, M.H. (2019). Dynamic REM Sleep Modulation by Ambient Temperature and the Critical Role of the Melanin-Concentrating Hormone System. Current Biology 29, 1976–1987.e4.

26. Scammell, T.E., Willie, J.T., Guilleminault, C., Siegel, J.M., and International Working Group on Rodent Models of Narcolepsy (2009). A consensus definition of cataplexy in mouse models of narcolepsy. Sleep 32, 111–116.

27. Oishi, Y., Williams, R.H., Agostinelli, L., Arrigoni, E., Fuller, P.M., Mochizuki, T., Saper, C.B., and Scammell, T.E. (2013). Role of the Medial Prefrontal Cortex in Cataplexy. Journal of Neuroscience 33, 9743–9751.

28. Szymusiak, R. (2018). Body temperature and sleep. Handb Clin Neurol 156, 341–351.

29. Heller, H.C. (2005). Temperature, thermoregulation and sleep. In Principles and Practice of Sleep Medicine (Elsivier Saunders).

30. Parmeggiani, P.L. (2003). Thermoregulation and sleep. Front. Biosci. 8, s557–567.

31. Cerri, M., Luppi, M., Tupone, D., Zamboni, G., and Amici, R. (2017). REM Sleep and Endothermy: Potential Sites and Mechanism of a Reciprocal Interference. Front Physiol 8, 624.

32. Harding, E.C., Yu, X., Miao, A., Andrews, N., Ma, Y., Ye, Z., Lignos, L., Miracca, G., Ba, W., Yustos, R., et al. (2018). A Neuronal Hub Binding Sleep Initiation and Body Cooling in Response to a Warm External Stimulus. Curr. Biol. 28, 2263–2273.e4.

33. Cerri, M., Ocampo-Garces, A., Amici, R., Baracchi, F., Capitani, P., Jones, C.A., Luppi, M., Perez, E., Parmeggiani, P.L., and Zamboni, G. (2005). Cold exposure and sleep in the rat: effects on sleep architecture and the electroencephalogram. Sleep 28, 694–705.

34. Szymusiak, R., Satinoff, E., Schallert, T., and Whishaw, I.Q. (1980). Brief skin temperature changes towards thermoneutrality trigger REM sleep in rats. Physiol. Behav. 25, 305–311.

35. Szymusiak, R., and Satinoff, E. (1981). Maximal REM sleep time defines a narrower thermoneutral zone than does minimal metabolic rate. Physiol. Behav. 26, 687–690.

36. Blumberg, M.S., Lesku, J.A., Libourel, P.-A., Schmidt, M.H., and Rattenborg, N.C. (2020). What Is REM Sleep? Current Biology 30, R38–R49.

37. Lo Martire, V., Silvani, A., Bastianini, S., Berteotti, C., and Zoccoli, G. (2012). Effects of ambient temperature on sleep and cardiovascular regulation in mice: the role of hypocretin/orexin neurons. PLoS ONE 7, e47032.

38. Tupone, D., Madden, C.J., Cano, G., and Morrison, S.F. (2011). An Orexinergic Projection from Perifornical Hypothalamus to Raphe Pallidus Increases Rat Brown Adipose Tissue Thermogenesis. Journal of Neuroscience 31, 15944–15955.

39. Sellayah, D., Bharaj, P., and Sikder, D. (2011). Orexin Is Required for Brown Adipose Tissue Development, Differentiation, and Function. Cell Metabolism 14, 478–490.

40. Imperatore, R., Palomba, L., and Cristino, L. (2017). Role of Orexin-A in Hypertension and Obesity. Curr Hypertens Rep 19, 34.

41. Sieminski, M., Szypenbejl, J., and Partinen, E. (2018). Orexins, Sleep, and Blood Pressure. Curr Hypertens Rep 20, 79.

42. Silvani, A., and Dampney, R.A.L. (2013). Central control of cardiovascular function during sleep. American Journal of Physiology-Heart and Circulatory Physiology 305, H1683–H1692.

43. Fronczek, R., Overeem, S., Lammers, G.J., van Dijk, J.G., and Van Someren, E.J.W. (2006). Altered skin-temperature regulation in narcolepsy relates to sleep propensity. Sleep 29, 1444–1449.

44. van der Heide, A., Werth, E., Donjacour, C.E.H.M., Reijntjes, R.H.A.M., Lammers, G.J., Van Someren, E.J.W., Baumann, C.R., and Fronczek, R. (2016). Core Body and Skin Temperature in Type 1 Narcolepsy in Daily Life; Effects of Sodium Oxybate and Prediction of Sleep Attacks. Sleep 39, 1941–1949.

45. Fronczek, R., Raymann, R.J.E.M., Overeem, S., Romeijn, N., van Dijk, J.G., Lammers, G.J., and Van Someren, E.J.W. (2008). Manipulation of skin temperature improves nocturnal sleep in narcolepsy. J. Neurol. Neurosurg. Psychiatr. 79, 1354–1357.

46. Fronczek, R., Raymann, R.J.E.M., Romeijn, N., Overeem, S., Fischer, M., Dijk, J.G. van, Jan Lammers, G., and Van Someren, E.J.W. (2008). Manipulation of Core Body and Skin Temperature Improves Vigilance and Maintenance of Wakefulness in Narcolepsy. Sleep 31, 233–240.

47. Raymann, R.J.E.M., Swaab, D.F., and Van Someren, E.J.W. (2008). Skin deep: enhanced sleep depth by cutaneous temperature manipulation. Brain 131, 500–513.

48. Raymann, R.J.E.M., Swaab, D.F., and Van Someren, E.J.W. (2005). Cutaneous warming promotes sleep onset. Am. J. Physiol. Regul. Integr. Comp. Physiol. 288, R1589–1597.

49. Kräuchi, K., Cajochen, C., Werth, E., and Wirz-Justice, A. (1999). Warm feet promote the rapid onset of sleep. Nature 401, 36–37.

50. Latifi, B., Adamantidis, A., Bassetti, C., and Schmidt, M.H. (2018). Sleep-Wake Cycling and Energy Conservation: Role of Hypocretin and the Lateral Hypothalamus in Dynamic State-Dependent Resource Optimization. Front Neurol 9, 790.

51. Aldrich, M.S., Chervin, R.D., and Malow, B.A. (1997). Value of the multiple sleep latency test (MSLT) for the diagnosis of narcolepsy. Sleep 20, 620–629.

52. Trotti, L.M., Staab, B.A., and Rye, D.B. (2013). Test-Retest Reliability of the Multiple Sleep Latency Test in Narcolepsy without Cataplexy and Idiopathic Hypersomnia. Journal of Clinical Sleep Medicine 09, 789–795.

53. Ruoff, C., Pizza, F., Trotti, L.M., Sonka, K., Vandi, S., Cheung, J., Pinto, S., Einen, M., Simakajornboon, N., Han, F., et al. (2018). The MSLT is Repeatable in Narcolepsy Type 1 But Not Narcolepsy Type 2: A Retrospective Patient Study. Journal of Clinical Sleep Medicine 14, 65–74.

54. Billiard, M. (2013). Multiple sleep latency test: are the practice and interpretation of the test valid? Sleep Med. 14, 127–128.

55. Lammers, G.J., Bassetti, C.L.A., Dolenc-Groselj, L., Jennum, P.J., Kallweit, U., Khatami, R., Lecendreux, M., Manconi, M., Mayer, G., Partinen, M., et al. (2020). Diagnosis of central disorders of hypersomnolence: A reappraisal by European experts. Sleep Medicine Reviews 52, 101306.

56. Baumann, C.R., Mignot, E., Lammers, G.J., Overeem, S., Arnulf, I., Rye, D., Dauvilliers, Y., Honda, M., Owens, J.A., Plazzi, G., et al. (2014). Challenges in diagnosing narcolepsy without cataplexy: a consensus statement. Sleep 37, 1035–1042.

