## Supplemental Figures for "Narcolepsy and the Dissociation of REM Sleep and Cataplexy through Ambient Temperature Manipulation"

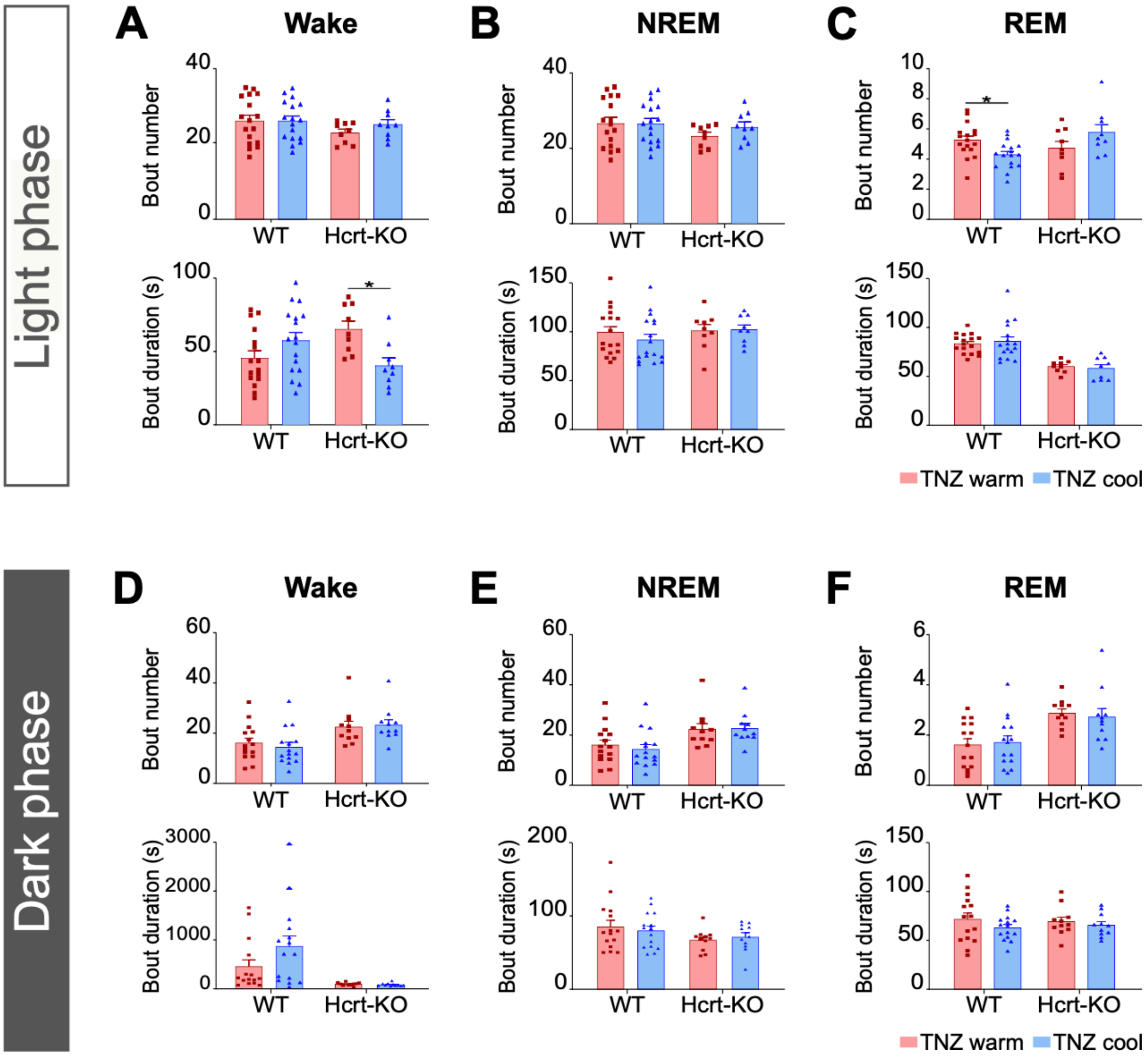


<u>**Supplementary Figure 1**</u>**. Subanalyses of sleep-wake expression in both the circadian light and dark phases. A-C)** Mean bout number and mean bout durations for wake, NREM sleep and REM sleep comparing the TNZ warm and TNZ cool conditions. **D-F)** Mean bout number and mean bout durations for wake, NREM sleep and REM sleep comparing the TNZ warm and TNZ cool conditions. Data were analyzed using two-way ANOVA and post-hoc Sidak's comparison test. Data are presented as means ± standard error of the mean (SEM). $^*p<0.05$.

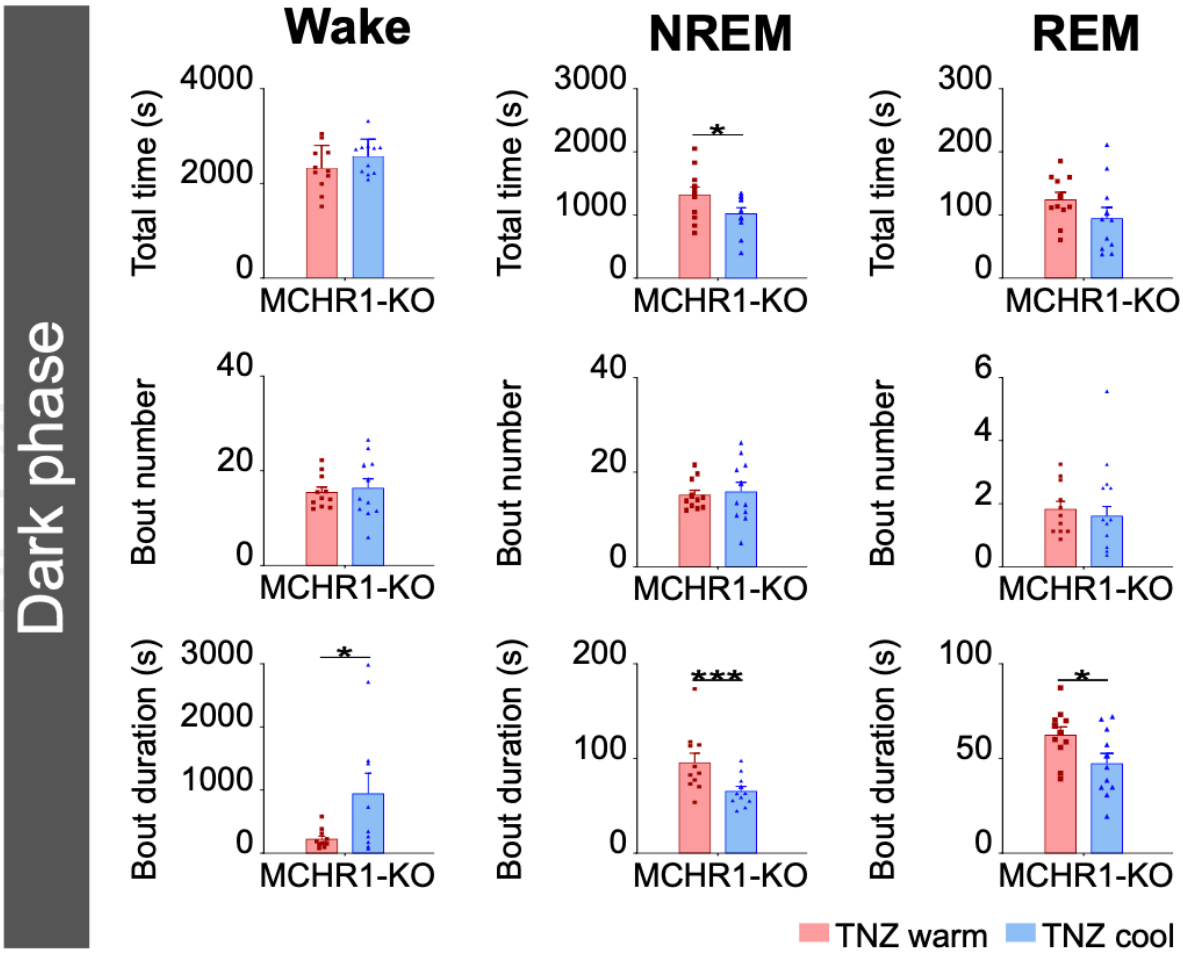


**<u>Supplementary Figure 2</u>. Sleep-wake expression for MCHR1-KO mice during the dark phase.** Data are presented as the total mean durations of wake, NREM sleep and REM sleep, as well as bout number and bout durations for each sleep state comparing the TNZ warm vs TNZ cool conditions. Data were analyzed using a two-tailed student's t test. Although there was a small but significant increase in NREM sleep during the TNZ warm condition, no significant changes were observed for mean total wake or REM sleep durations as a function of Ta condition. Data are presented as means ± standard error of the mean (SEM). *$p<0.05$; ***$p<0.001$.

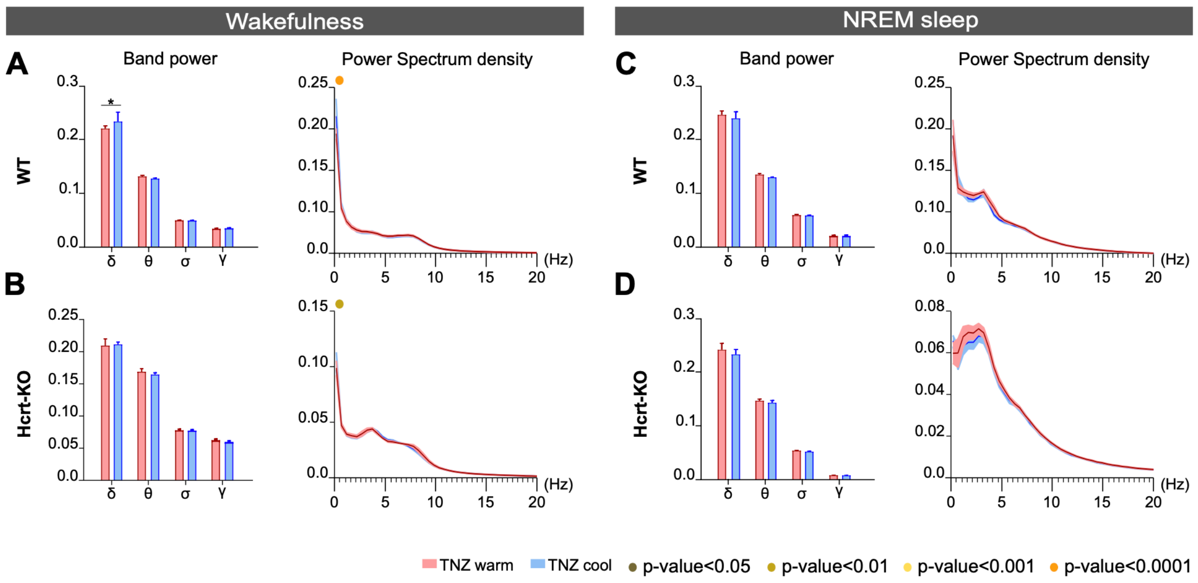


<u>Supplementary Figure 3</u>. **EEG Band power and power spectral density analyses for wakefulness (A-B) and NREM sleep (C-D) for WT and Hcrt-KO mice.** Other than an isolated increase in the delta band for the WT mice during the Ta cooling phase, no other significant differences were observed as a function of Ta condition.
